# Vascular plant extinctions outpace colonizations under climate change

**DOI:** 10.64898/2025.12.26.696646

**Authors:** Barnabas H. Daru, Siri Birkeland, Desalegn Chala

**Affiliations:** Department of Biology, Stanford University, Stanford, CA 94305, USA; Natural History Museum, University of Oslo, 0562 Oslo, Norway

**Author notes:** Barnabas H. Daru, **Email:**.

## Abstract

Climate change is expected to shift plant distributions poleward and upslope, yet requires global empirical evidence. Using historical range shifts from global herbarium specimens for 109,242 vascular plant species over the past 28 years as baseline, we fit 1.8 million species distribution models on a global dataset of 184,545 vascular plant species to project future range shifts over an equivalent period, the next 20 years (2021-2040). We find that plant species have already shifted their distribution centroids at a median velocity of 5.6 km per year globally, exceeding projected future velocities of 3.27 km yr per year. Projected shifts will vary widely in direction and magnitude, with varied velocities, colonization and extinction rates. Shifts will vary across clades with Superasteridae, Gymnosperms and Monocots moving at slower velocities, and Ferns, Chloranthales, and Magnoliidae with faster velocities. Although most clades will shift polewards, velocities will be highest in temperate and boreal regions, while shifts will be slower in regions of high topographic variation in the tropics. High latitude species will trend poleward and upslope, while desert shrubs, and tropical conifer forests will show slower and directional shifts. Extinctions will outpace colonizations, confirming that climate change will be a driver of biodiversity loss. Colonization will be frequent in protected areas, higher elevations, and among threatened species whereas extinctions will cluster near species’ historical ranges and at lower elevations. Our findings show the complexity of climate-driven range shifts and emphasize the importance of considering multiple interacting processes beyond simple latitudinal or elevational movements.

## Main Text

Climate change is a major driver of biodiversity change and is expected to alter species distributions in the future (*1–3*). As climate changes, species may respond by shifting their ranges to track suitable habitats, adjust through physiological plasticity, evolve to tolerate new conditions, or go extinct due to suboptimal conditions (*4*). How prevalent each of these responses are across species and biomes remains poorly understood. However, range shifts have emerged as one of the most well-documented responses to climate change, with species moving across latitude, longitude and altitude to maintain suitable environmental conditions (*5*). These shifts can result in local extinctions as species fail to track changing climates, and colonization, where species establish in new areas, which can reshape spatial patterns of biodiversity and ecosystem functioning (*6, 7*). However, the velocity and extent of these shifts are still poorly understood, especially across diverse plant clades and geographic regions.

Studies investigating climate-driven range shifts in plant species have mostly been performed at local or regional scales in temperate regions of the northern hemisphere, have focused on poleward and upslope range shifts patterns (*2, 5, 8–10*), and have relied on long-term vegetation surveys. A key assumption is that climate (especially global temperatures) is correlated with latitude and altitude, and thus species are expected to track their specific thermal tolerance ranges at higher latitudes and altitudes in response to warming (*5, 11–13*). However, this approach overlooks plant range shifts in biodiversity-rich tropics where many species are vulnerable to climate change due to their narrower climatic tolerances (*14*) and thus may experience more complex, multidimensional shifts (*15*). Given that tropical species are often ecological specialists with narrow climatic niches (*16*), we hypothesize that their responses to climate change will differ from those of generalist species in temperate regions, leading to more constrained or idiosyncratic range shifts. Similarly, species in regions with lower climatic seasonality, extreme environments like cacti and desert plants, or those near their upper thermal limits are more likely to experience negative responses to climate change (*17, 18*), which could disproportionately affect tropical floras. Likewise, while long-term vegetation plot datasets are increasingly becoming available and now cover wide spatial and temporal resolutions (e.g., https://forestreplot.ugent.be/ or https://euroveg.org/resurvey/), there are still large coverage gaps in other parts of the world (*19, 20*).

Millions of plant occurrence records from over 3,000 herbaria worldwide and citizen science programs can bridge geographic and temporal gaps by providing time series data to compare past, present, and projected ranges in response to climate change, land-use change, and other anthropogenic drivers (*21, 22*). To establish a historical baseline for ecological change in plant communities, we analyzed 16,219,436 herbarium records representing 109,242 vascular plant species across the globe in their native ranges, spanning two recent periods of climate change (1970–2000, median 1983, and 2001–2024, median 2010). Over the past 28 years, plant species have shifted the center of their distributions (centroids) at a median (±s.e.m.) velocity of 5.6±0.06 km/yr (95% CI for median: 5.5 to 5.6 km/yr), with a tendency for high latitude regions to shift poleward (Rayleigh’s *r* = 0.06, *P* < 0.05; Fig. 1). If this trend represents realized change that has happened already over the past 28 years, then observed historical range shifts provide an important empirical benchmark for evaluating future projections. We test these expectations by predicting plant range shifts under current and future climate change at a global scale.

**Figure 1.**
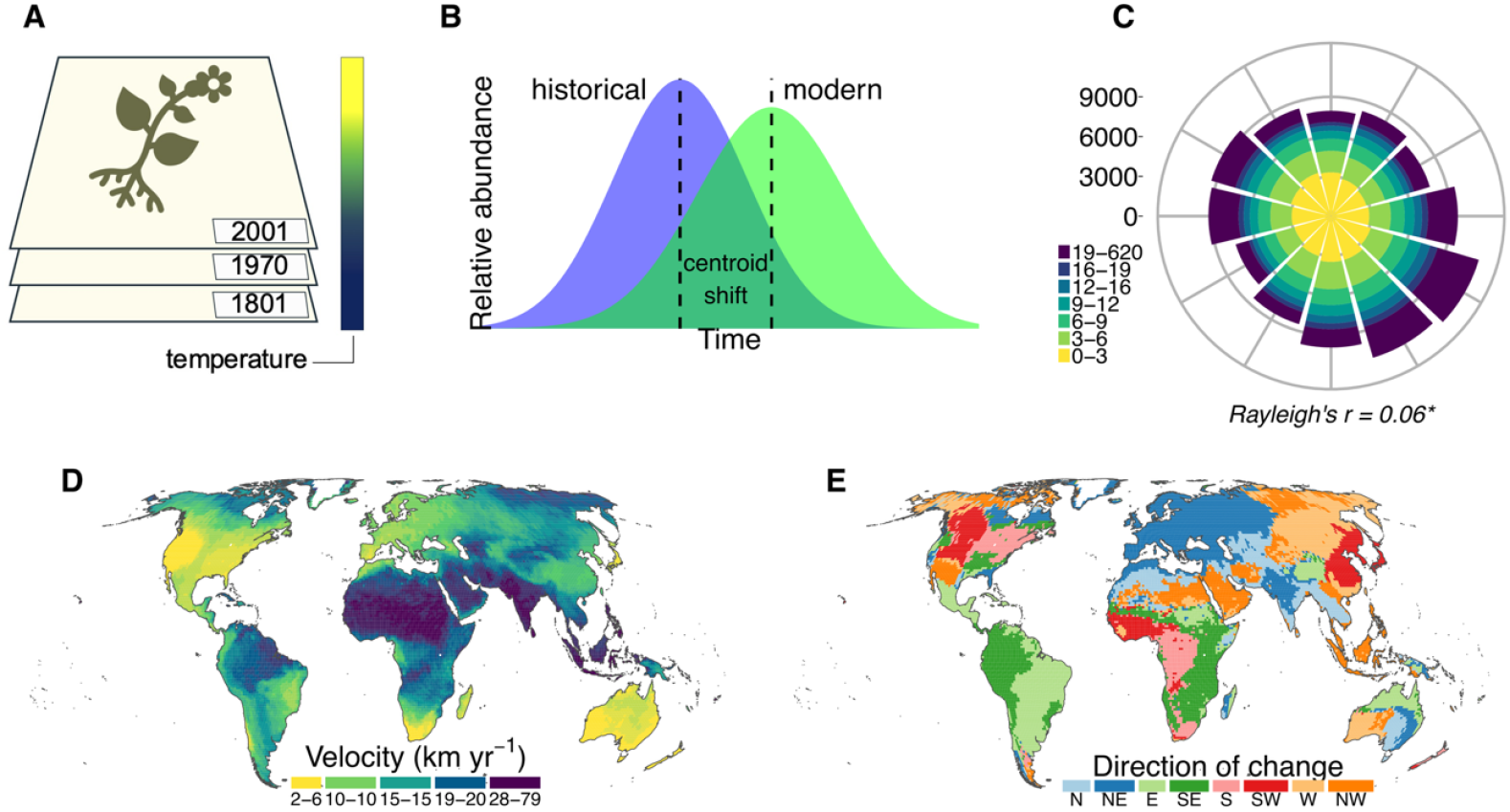
Baseline shifts in plant distributions across the globe in the past from 1970 to 2024 based on herbarium records. **A**, Piles of herbarium specimens to show the temporal breadth captured by herbarium collections. **B**, Hypothesized centroid shifts in species distribution centers between historical (blue) and modern (green) ranges. **C**, Wind rose diagram showing the directional tendency and velocity of species’ centroid shifts between the two time periods. **D**, Geographic variation in range shift velocity, with colors indicating different magnitudes of movement. **E**, Geographic variation in compass bearing of range shifts showing trends in plant distributions in the past 28 years.

Apart from climate change, future plant range shifts may be shaped by other factors including intrinsic functional traits, biotic interactions, abiotic conditions, and other anthropogenic pressures. High dispersal (e.g., small seeds) may drive faster shifts, while conservative traits slow shifts or raise extinction risk. Plant interaction with other organisms such as insects can mediate how changing climates influence range shifts, and these effects are expected to vary across regions (*23*). Abiotic factors such as climate variability, topographic heterogeneity, and environmental extremes can influence both the velocity and direction of shifts, of which we predict faster rates in climatically unstable regions and slower and more multidirectional shifts in rugged terrain with steep gradients and microrefugia. Finally, we predict that anthropogenic pressures are expected to influence plant movement, colonization potential, and local extinction risk, particularly in biodiversity-rich but highly modified landscapes. For instance, land-use conversion and fragmentation will reduce connectivity (*24*) by slowing movement and increasing trailing-edge extirpations whereas protected areas and intact elevational corridors will show higher colonization and persistence (*25, 26*) because of continuous habitat connectivity and reduced edge effects and human disturbance. These expectations provide a framework for predicting how biotic and abiotic factors interact to shape future plant distributions under climate change.

Here, we use species distribution models (SDMs) to project potential suitable habitat for 184,545 vascular plant species over the next 20 years under nine future global climate models and scenarios between 2021 and 2040. Our analysis incorporate sampling bias correction, evolutionary dispersal rates based on phylogeographic spherical Brownian motion model (*27*), uncertainty across multiple global climate models and scenarios (*28–30*), and a suite of ecological predictors and biotic interactions of butterflies within an SDM framework to enable us capture the complex processes that shape range shifts across the globe, clades, and biomes. Specifically, we address three questions: (1) How will climate change affect range shifts across clades and regions under future climate change scenarios? (2) What factors predict projected plant range shifts at a global scale? (3) How do the predictors of projected velocity shift, colonization, and local extinction differ?

## Results and Discussion

### Species-level range shift velocities based on future modelled projections

We modelled climate niche shift estimates based on an ensemble of three global climate models: warm (UKESM1-0-LL) (*29*), dry (MPI-ESM1-2-HR) (*28*), and wet (IPSL-CM6A-LR) (*30*) under three greenhouse gas emission scenarios, optimistic (SSP126), high (SSP370), and extreme (SSP585) for a total of 1,845,450 SDMs. We project that over the next 20 years (between 2021 and 2040) plants will shift their centroids at a median (±s.e.m.) velocity of 3.27± 0.063 km yr^−1^ based on the centroid of current and projected distribution maps (25^th^ to 75^th^ percentiles = 1.78 to 6.35 km yr^−1^, *n* = 184,549 species). By contrast, global historical estimates derived from herbarium specimens indicate that plant species have already shifted their distribution centroids at a higher median velocity of 5.6 km yr□^1^ over the past 28 years (Wilcoxon rank test, one-sided: P < 2.2 × 10^−16^). This result indicates that plant distributions are already reorganizing at velocities comparable to and even exceeding those projected for the coming decades. Throughout, we summarize uncertainty in our results by reporting the median estimates of range shifts across the nine climate scenarios.

The shift rates are projected to be phylogenetically patterned such that among major plant clades, Superasteridae, Gymnosperms, and Monocots will experience the slowest median velocity shifts of 3.03±0.093 km yr^−1^, 3.08±0.38 km yr^−1^, and 3.11±0.14 km yr^−1^ respectively. This may indicate heightened vulnerability to climate change due to potential limitations in tracking suitable environmental conditions, which could elevate extinction risks (31). Conversely, Ferns, Chloranthales and Magnoliidae will experience the fastest velocity shifts of 4.86±0.53 km yr^−1^, 4.65±5.46 km yr^−1^, and 3.76±0.30 km yr^−1^, respectively (Fig. 2), likely due to traits like superior dispersal and rapid generation times in these clades (*3*). Beyond speed, there will be differences in the compass bearing of range shifts (Fig. 2). Most clades will trend northwestward including Superasteridae (327°), Basal Eudicots (286°), and Superrosidae (320°) while Monocots (12°) and Ferns (8°) are projected to trend northeastward. Directionality is projected to be weak overall (Rayleigh’s *r* = 0.03–0.12), most likely because of broad but clade-specific compass bearings rather than a single global axis of movement. Thus, our multidimensional range shift analyses reveals that vascular plants will experience shifts in all directions, at different velocities across clades.

**Figure 2.**
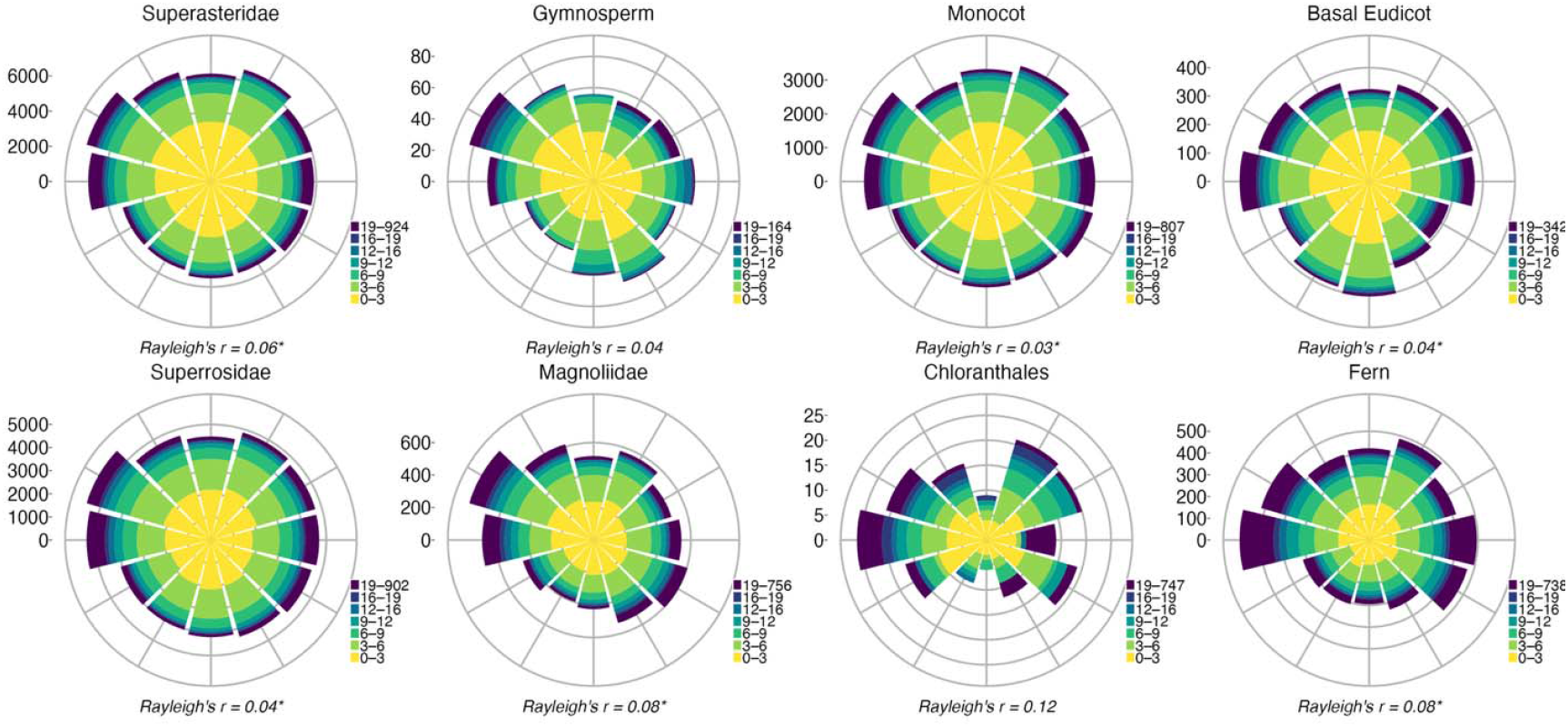
Projected velocity and direction of range shifts for major vascular plant lineages (*n* = 184,545 species) between current and future climate change in 2021-2040. Each polar plot (wind rose) summarizes the projected directional and velocity distributions for each plant lineage. The radial axis indicates the number of species showing shifts in a given direction, while the colored rings indicate the range of velocity of range shifts (in km year□^1^). Warmer colors (yellow) represent slower shifts and cooler colors (purple) indicate more rapid shifts. The Rayleigh’s *r* statistic below each plot quantifies the strength of the directional trend in niche shifts (^*^ *P* < 0.05 indicates significant directionality). The wind-roses are plotted based on an ensemble of three global climate models: warm (UKESM1-0-LL), dry (MPI-ESM1-2-HR), and wet IPSL-CM6A-LR) and each for three greenhouse gas emission scenarios, optimistic (SSP126), high (SSP370), and extreme (SSP585), resulting in a total of nine climate scenarios. Wind-roses were created with custom R scripts from ref.(39).

### Geographic patterns of projected range shift velocities

#### Projected range shift velocity

Range shifts velocities will vary substantially among clades and regions, and these patterns will not be geographically uniform across latitudinal gradients (Moran’s *I* = 0.15, *P* ≪ 0.001, Fig. 3). While the magnitude depends on the climate models and emission scenarios used, the spatial trends are consistent across scenarios. The fastest shifts are projected in high-latitude and lowland ecosystems of the Northern Hemisphere, northern North America, northern Europe, and northern Asia, where velocities gradually decline toward the equator and Southern Hemisphere (Fig. 3a, b). This pattern is consistent with the baseline evolutionary dispersal rate of species based on phylogeographic spherical Brownian motion model (SBM) (*27*) which reflects biologically informed constraints on species’ potential movements, i.e., constraints based on species evolutionary history before any modern anthropogenic change (27, 32, 33). Consequently, regions that have historically experienced high evolutionary dispersal rates over millions of years like the northern temperate zones of North America and Eurasia, are projected to continue experiencing high range shift velocities in the coming decades (Pearson’s *r* = 0.52 for SBM dispersal rates vs. projected velocities; *P* ≪ 0.0001). By contrast, areas with high topographic variation especially in tropical and subtropical regions, that is, southern North America, Neotropics, Afrotropics, Southeast Asia, and Australasia, will experience slower shift velocities (Fig. 3a). Most of these areas overlap with high elevation regions like Andes, Hengduan-Himalaya, or Eastern Arc of East Africa, and harbor high plant diversity (34, 35) which are projected to face some of the slowest velocities of climate change globally (36). This disparity suggests that plant taxa may be unable to move quickly enough to track shifting climatic conditions, leading to climate-driven range mismatches, a pattern already observed in some taxonomic groups (37). These projected poleward and elevational shifts are consistent with broader patterns of climate-driven range shifts documented in various taxonomic groups (3, 5, 38).

**Figure 3.**
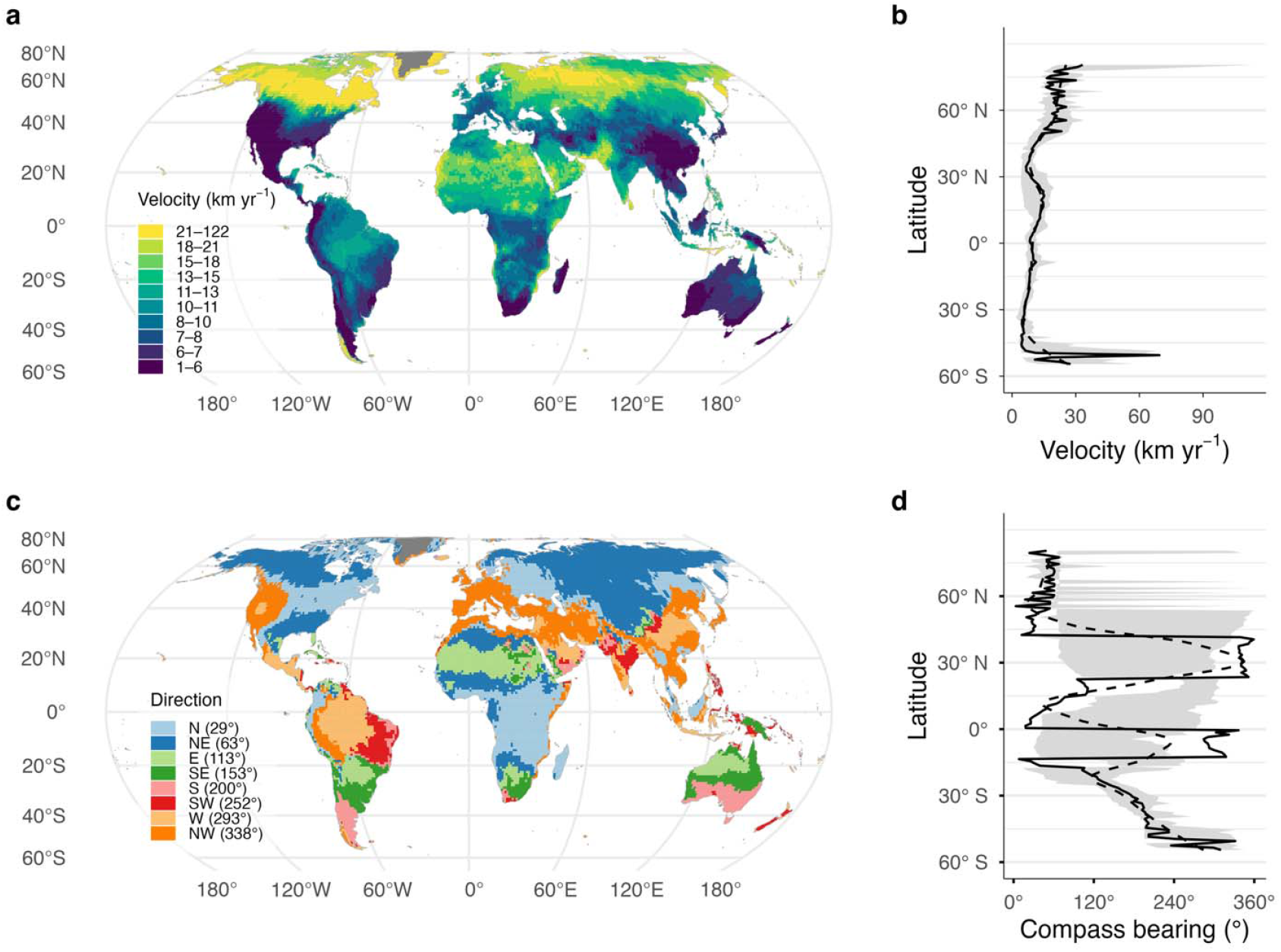
Global patterns of projected species range shift velocity, compass direction and latitudinal trends under future climate change in 2021-2040. **a**, Magnitude of projected velocity of range shift (km yr^−1^) calculated as the median rate at which a species environmental niche is projected to move across grid cells in 20 years. Dark purple colors indicate slower shifts, and yellow colors indicate faster shifts. **b**, Latitudinal trend of median range-shift velocity; shaded ribbon shows the range (10^th^ to 90^th^ percentiles) across 1° latitude bands; dashed line is a LOESS smooth of the median. **c**, The projected compass bearing of range shift velocities, expressed in degrees relative to north. Colors indicate the compass bearing (N, NE, E, SE, S, SW, W, NW) of range shift displacement pathways. **d**, Latitudinal trend of median compass bearing; ribbon = 10^th^ to 90^th^ percentiles across 1° bands; dashed line = circular LOESS smooth. **a, c**, Maps are shown at a 100 km × 100 km spatial resolution. Areas masked in grey indicate sparsely vegetated regions based on global land-cover data. The maps are plotted based on an ensemble of three global climate models: warm (UKESM1-0-LL), dry (MPI-ESM1-2-HR), and wet IPSL-CM6A-LR) and each for three greenhouse gas emission scenarios, optimistic (SSP126), high (SSP370), and extreme (SSP585), resulting in a total of nine climate scenarios. The maps are in the Equal-Earth projection system.

#### Projected shifts across compass bearing

When considering the compass bearing of range shifts normalized to 0-360°, we classified northward shifts as poleward and southward as equatorward in the Northern Hemisphere, with the opposite in the Southern Hemisphere (that is, southward as poleward and northward as equatorward). We found a tendency for high latitude regions to shift poleward (that is, northward in the Northern Hemisphere and southward in the Southern Hemisphere) (Fig. 3c,d). Species in northern North America, and Eurasia are expected to experience a net northward to northeastward shift (Fig. 3c). By contrast, southward to southeastward shifts are anticipated in the Amazon, Patagonia, Cape Floristic Region, India, southern Australia and New Zealand. The poleward shifts in high-latitude regions likely reflect species tracking isotherms under warming (*3, 5*). However, several areas also show strong longitudinal shifts: eastward shifts are projected in arid regions such as subtropical South America to Atacama deserts, the Sahara, northern Southern Africa, and the Eremean region of Australia, whereas westward shifts are projected in the southwestern United States, Atlantic Forest, and the Mediterranean through the Middle East, and East Asia (Fig. 3c). We also tested whether our results were sensitive to the choice of climate model or emission scenario (SI Appendix, Fig. S3). While variations in climate models and scenarios altered directionality in regions like Amazonia, Australasia, and East Asia, trends remained similar, and support our key finding that climate change will alter plant distributions in multiple directions. These patterns indicate that climate-driven range shifts are not only latitudinal but reflect regional climatic and environmental gradients as already found for some regional floras (39).

#### Projected shift across biomes

Across biomes of the world (40), the fastest median centroid shifts are projected to occur in tropical grassland and shrublands (7.1 km yr^−1^; NE, 59°), tropical dry forest (5.6 km yr^−1^; NE, 46°) and mangrove (5.3 km yr^−1^; E, 95°). By contrast, desert shrublands and tropical conifer forests are projected to shift the slowest (both 2.0 km yr□^1^). The remaining biomes are expected to show intermediate velocities and bearings from northward to northwestward (Fig. 4). The concentration of range shifts in a single direction is projected to be very weak especially in Mediterranean biomes, and only slightly stronger in desert shrub and montane grasslands, which means that species in these biomes will move in multiple directions rather than predominantly poleward (Rayleigh’s *r* = 0.02, 0.14, and 0.07 respectively; all *P* < 0.05). Our findings of range shift velocities across biomes are robust to varying global climate models and to scenarios of greenhouse gas emission.

**Figure 4.**
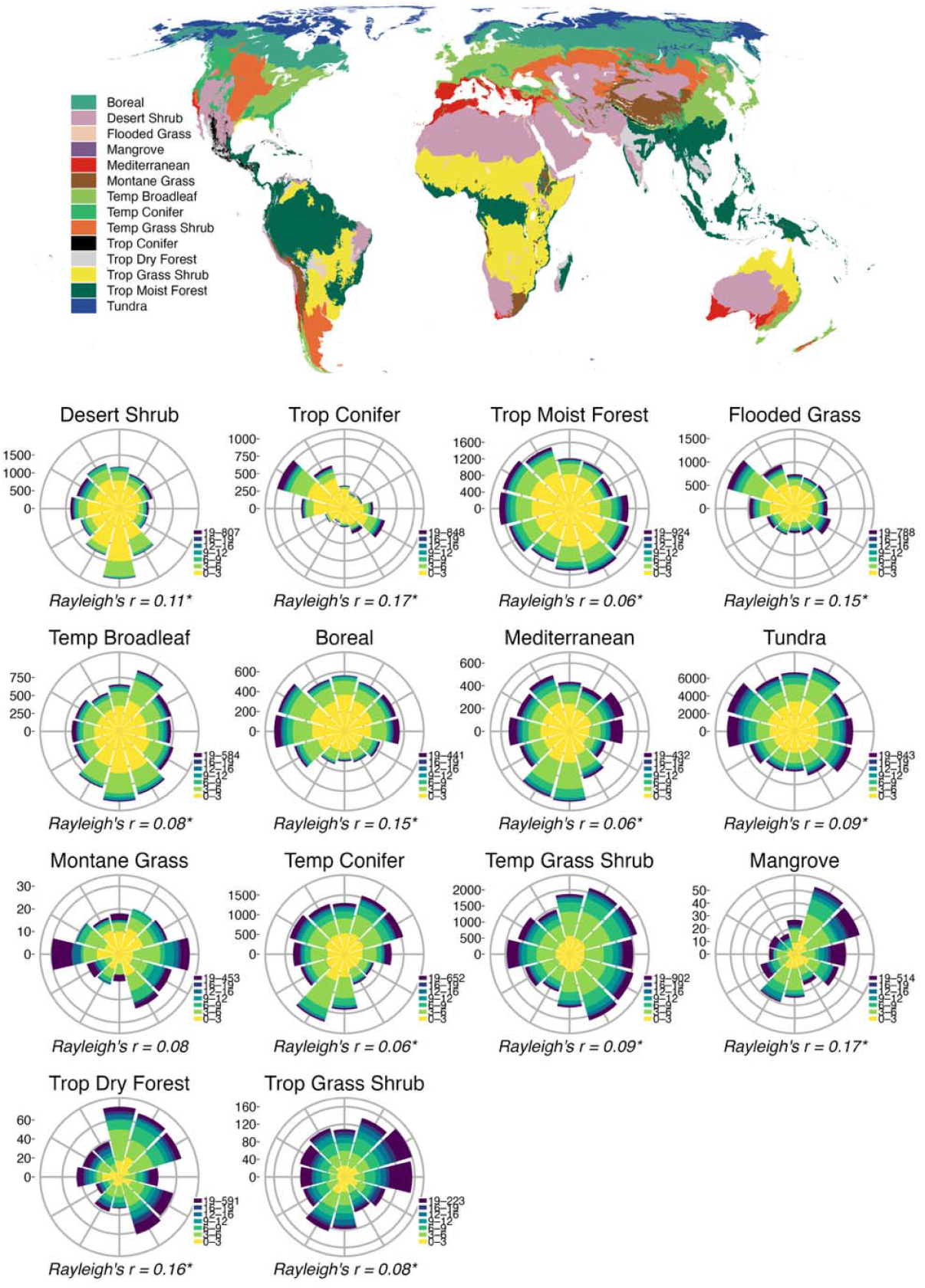
Projected velocity and direction of species’ centroid shifts across biomes of the world. Top, the distribution of terrestrial biomes shown on an Equal-Earth projection (EPSG:8857). Bottom, wind-rose plots summarizing the direction and velocity of centroid shifts for species occurring in each biome. Bars indicate the number of species shifting in each 30° sector, and color encodes shift velocity (km yr^−1^) in fixed discrete bins. Legend tick labels show range values of velocities. Rayleigh’s *r* indicates directional concentration for each biome (asterisk denotes *P* < 0.05). The wind-roses are generated based on an ensemble of three global climate models: warm (UKESM1-0-LL), dry (MPI-ESM1-2-HR), and wet IPSL-CM6A-LR) and each for three greenhouse gas emission scenarios, optimistic (SSP126), high (SSP370), and extreme (SSP585). Wind-roses in bottom panels were created with custom R scripts from ref (39). Ecoregions are based on Terrestrial Ecoregions of the World(40), and the world basemap was derived from shapefiles available at https://naturalearthdata.com.

#### Projected colonization versus extinction

We identify areas of projected range loss and gain under future climate change because colonization at the leading edge and extinction at the trailing edge are driven by different mechanisms (41). This spatial decoupling can create extinction debts and colonization credits (42). To this end, we compared centroid shifts for each species associated with colonization (that is, new climate niches where species are moving into) versus extinction (where climatic suitability declines) based on the ensembled model (3 GCMs × 3 SSP scenario). Across species, the median difference was -12.95 percentage points (95% CI: -13.08 to -12.82), which indicates that extinctions predominate over colonizations (Wilcoxon signed□rank test: *V* = 4.4 × 10^9^, *P* ≪ 0.001). This finding means that species will lose climatically suitable habitats faster than they can establish themselves in new ones, likely due to asymmetric climate change effects, dispersal limitations, or habitat fragmentation restricting successful range expansions (43–46). Previous work has shown that non-native naturalizations and colonizations promote biotic homogenization of plant communities in the Anthropocene (47, 48). Our finding that extinctions will outpace colonizations adds a further layer of concern, which means that the combined pressures of non-native naturalizations and projected local extinctions act as forces to reshape plant communities in the Anthropocene. The hotspots of local extinctions are projected in biodiversity-rich regions like Amazonia, Central Africa, Southeast Asia, the Mediterranean, and parts of Australasia (Fig. 5e), where high endemism and anthropogenic pressures elevate extinction risks (49). Colonization hotspots will be less frequent but also projected in those same areas, especially in some parts of South America and East Asia (Fig. 5e).

**Figure 5.**
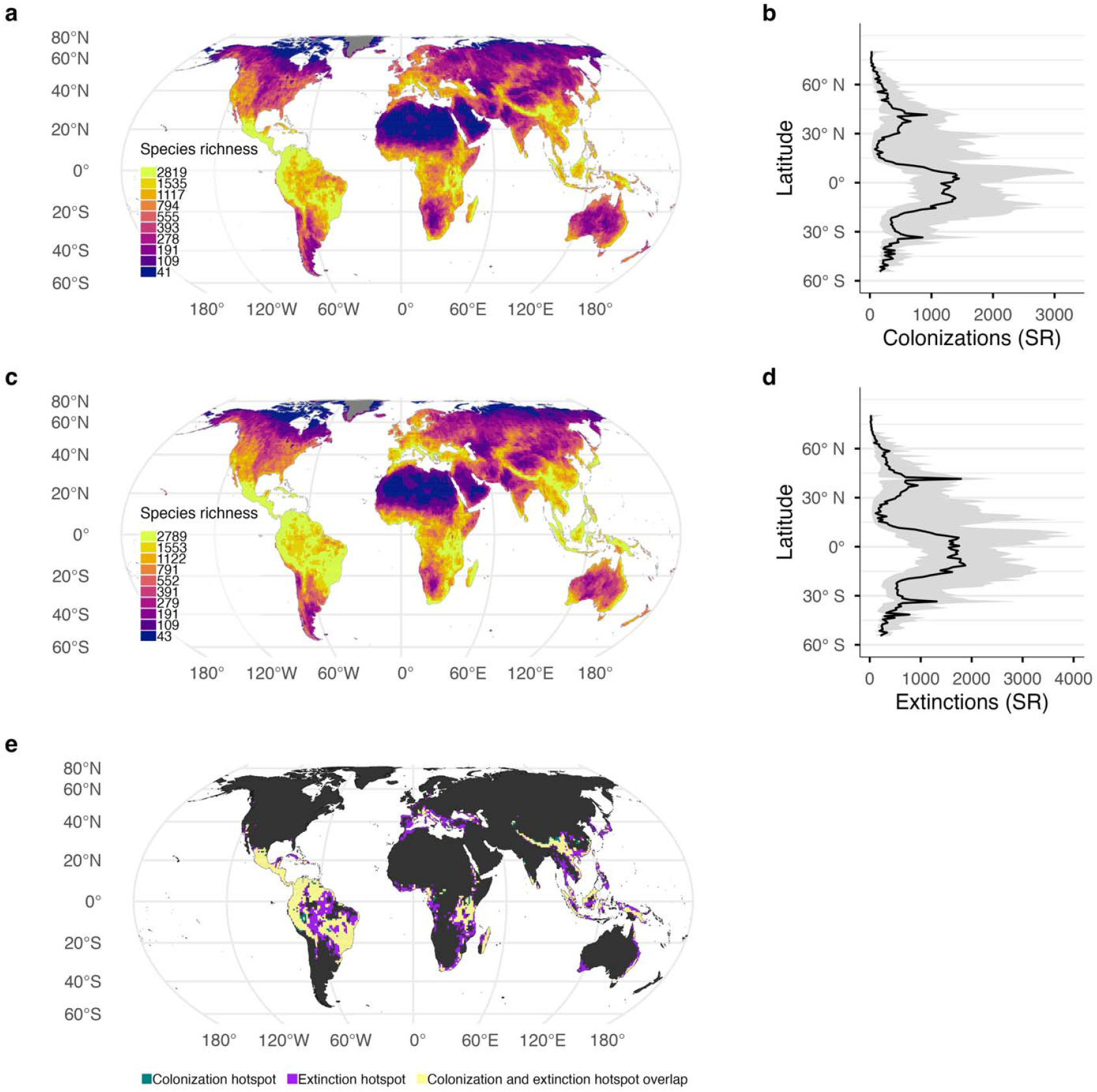
Global patterns and latitudinal trends of projected local colonizations and extinctions of vascular plant range shifts under future climate change. **a**, Total number of colonizations defined as the number of new climate niches where species are moving into, aggregated across 100 km grid cells, **b**, Latitudinal trend of projected median colonization; shaded ribbon shows the interdecile range (10^th^ to 90^th^ percentiles) across 1° latitude bands; dashed line is a LOESS smooth of the median. **c**, Total number of projected local extinctions defined as 100 km grid cells where species’ climatic suitability is projected to decline, **d**, Latitudinal trend of projected median extinction; shaded ribbon shows the interdecile range (10^th^ to 90^th^ percentiles) across 1° latitude bands; dashed line is a LOESS smooth of the median, and **e**, Hotspots defined as the top 10% richest cells in the proportion of species range shifts that are projected to become locally extinct (purple) or colonized (teal) under future climate change. Areas where extinctions and colonization overlap are indicated in yellow. The maps are plotted based on an ensemble of three global climate models: warm (UKESM1-0-LL), dry (MPI-ESM1-2-HR), and wet IPSL-CM6A-LR) and each for three greenhouse gas emission scenarios, optimistic (SSP126), high (SSP370), and extreme (SSP585), resulting in a total of nine climate scenarios. SR, species richness. The maps are in the Equal-Earth projection system.

### Predictors of projected species range shifts, colonization and local extinction

We used a linear mixed-effects models to analyze how ecological predictors will influence projected velocity shifts, local extinctions, and colonizations. We categorize these predictors into four broad groups: functional traits, biotic interactions, abiotic factors, and anthropogenic pressures (Fig. 6), including random intercepts for plant family and biome. Because SDM projections are driven mainly by climate, we interpret abiotic predictors as correlates of spatial variation in projected climatic displacement, and anthropogenic, biotic, and trait predictors as baseline proxies for potential constraints or facilitators of future realized tracking, not as causal drivers of the projections.

**Figure 6.**
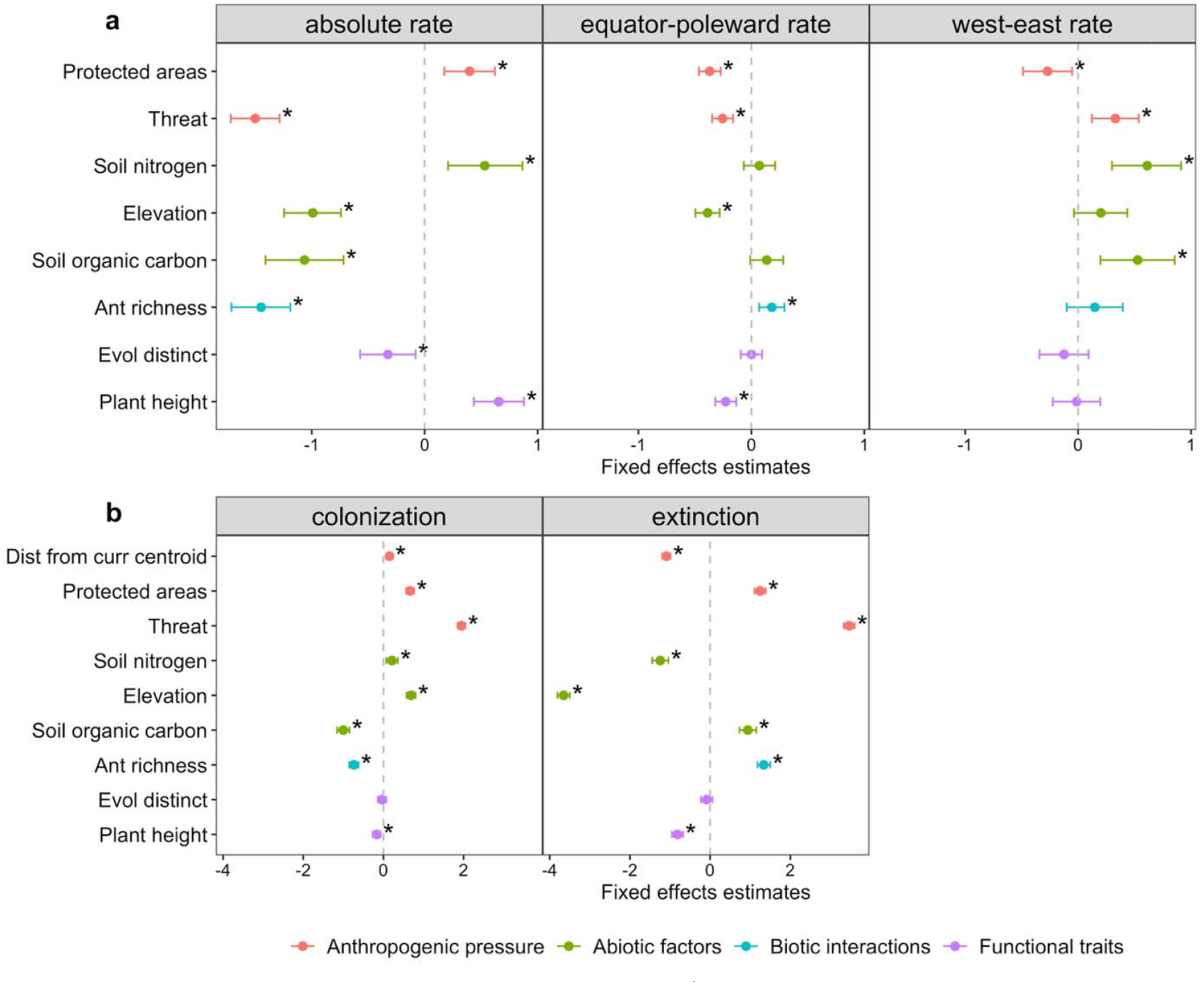
Determinants of projected range shift velocity (km yr^−1^), colonization, and extinction. **a**, Standardized fixed effects (±□ 95% CI) from a linear mixed-effects model of the effect of anthropogenic, biotic, abiotic, and functional trait predictors on absolute shift rates (left), equator-poleward shifts rates (middle) and west-east shift rates (right). **b**, Estimate effects of predictor variables on local colonization (left) and extinction events (right), where colonization refers to the successful establishment of a species in a previously unoccupied location, and extinction refers to the disappearance of a species from a site where it previously occurred. Each response variable was derived from an ensembled model of three GCMs × three SSP scenarios. Each predictor is color-coded according to its ecological grouping: pink for anthropogenic pressure, green for abiotic factors, teal for biotic interactions, and purple for plant functional traits. Points to the left of zero indicate a negative effect on shift velocities, whereas points to the right indicate a positive effect. The equator-poleward rate (km yr^−1^) indicates species’ projected latitudinal shifts, with positive values indicating poleward shifts and negative values equatorward shifts. The west-east rate (km yr^−1^) captures longitudinal movements, where positive values denote eastward and negative values westward shifts. Error bars represent 95% confidence intervals. All continuous predictors were z-scored; models include random intercepts for plant family and biome. Asterisks indicate predictors with P < 0.05 (two-sided tests).

We found that anthropogenic threat negatively impacts projected absolute and equator-poleward velocity shifts (β = -1.50 and -0.26, *P* ≪ 0.01; Fig. 6a), which indicates slower range shifts in threatened areas. Regions of higher soil nitrogen strongly associates with greater range movement and a tendency for west-east shifts (β = 0.53 and 0.61 respectively, *P* ≪ 0.01; Fig. 6a). This suggests that nitrogen deposition may accelerate range changes, a pattern already observed in European forest plants (39). Elevation and soil organic carbon negatively affect absolute velocity shifts. For biotic interactions, ant richness negatively predicts absolute velocities but positively predict equator□poleward and west□east shifts (Fig. 6a) most likely because climate□structured ant-plant mutualisms and short distance myrmecochory increase local retention while channeling movement along temperature and precipitation gradients (50). Finally, for intrinsic functional traits, plant height positively predicts absolute velocity shifts (β = 0.66, *P* ≪ 0.01) but negatively influence latitudinal and longitudinal shift velocities (Fig. 6a). Evolutionary distinctiveness negatively associates with absolute rate shifts meaning that evolutionarily distinct species (those with fewer or no close living relatives) will shift the slowest. These findings show the combined role of anthropogenic pressures, edaphic factors, biotic interactions, and functional traits in shaping plant isotherm shifts, with nutrient availability and biotic interactions facilitating movement, while topographic constraints, human threats, and evolutionary distinctiveness tend to slow plants down. This means that plant responses to future climate change will be complex and require ecological, environmental factors, and evolutionary drivers.

Colonization events will be more likely in protected areas, at higher elevations, and among threatened species (Fig. 6b). As species track shifting isotherms under anthropogenic climate warming (3, 51) and migrate toward more suitable habitats, these patterns indicate the role of conservation in facilitating climate-driven isotherm shifts. Protected areas act as stable refugia, which support natural colonization as species adjust to changing conditions (52), and suggest that protected areas could still be effective under a changing climate as already demonstrated for tetrapods (53). Elevated terrains provide cooler microclimates, consistent with documented upward shifts in species distributions where higher elevations serve as thermal refuges (54). In contrast, extinction events will be more common near species’ historical ranges, at lower elevations, and among smaller stature plants (Fig. 6b). These findings suggest that many species will be unable to adapt quickly enough to rapidly changing conditions, particularly in already disturbed or ecologically stressed areas (2, 55).

Altogether, our projections reveal variation in plant range shift velocities among clades: Ferns and Chloranthales are expected to move rapidly, while Superasteridae and Gymnosperms will shift more slowly and in less directional ways, likely reflecting dispersal limitations and habitat constraints. Geographically, temperate and boreal regions will experience the fastest shifts, consistent with the relationship between warming rates and species movement (36). By contrast, slow shifts in areas with high topographic variation especially in tropical and subtropical regions such as Andes, Hengduan-Himalaya, or Eastern Arc, may result from high species richness, habitat fragmentation, and complex biotic interactions. Isotherm shifts are expected to be strongly directional, with poleward movement dominating at high latitudes and longitudinal shifts most pronounced in southwestern United States, Amazonia, and southern Eurasia. At the biome level, desert ecosystems and tropical coniferous forests are projected to shift slowly, whereas tropical grasslands and shrublands are expected to move more rapidly, reflecting the influence of ecosystem structure. Our models indicate that isotherm shifts will be shaped, not only by climate but also by abiotic factors, functional traits, biotic interactions, and evolutionary history. Plant height and soil nitrogen emerged as strong predictors, whereas anthropogenic factors and topographic constraints slowed movement. Importantly, local extinctions will outpace colonization, suggesting that species will lose suitable habitats faster than they can establish in new areas. This pattern is especially pronounced in biodiversity-rich regions like Amazonia, Central Africa, Southeast Asia, and Australasia where high endemism and anthropogenic pressures may increase extinction risks. Collectively, these findings demonstrate that plant responses to future environmental change will be complex, varying across climatic, ecological, and evolutionary contexts worldwide.

## Materials and Methods

We compiled 498 million GBIF plant occurrences, reconciled names with the World Checklist of Vascular Plants, restricted records to native TDWG regions, removed introduced/doubtful points, spatially thinned occurrence records at 5 arcmin resolution, and constructed alpha-hull range polygons (10 km buffers for ≤3 points) from which we drew pseudo-occurrences scaled by range size. Dispersal constraints were incorporated using clade-level evolutionary dispersal rates estimated from a phylogeographic spherical Brownian motion model (27, 32) fitted to a global dated phylogeny(56), which we used to define species-specific training areas intersected with terrestrial ecoregions (40). Background points were sampled probabilistically from a kernel-density bias surface and also incorporated in our species distribution modeling. Environmental predictors (10 arcmin) included WorldClim v2.1 present (1970–2000) and nine future combinations (UKESM1-0-LL, MPI-ESM1-2-HR, IPSL-CM6A-LR × SSP126/370/585), plus butterfly richness to account for biotic interactions in our species distribution modeling; multicollinearity was reduced using variance inflation factors. We trained MaxEnt species distribution models with cross-validation and tuning, thresholded at the 95% occurrence quantile to produce binary maps, and generated 1,845,450 SDMs for 184,545 species (current + future). Performance was evaluated with AUC, TSS, and Boyce Index. For projections, we computed geodesic centroid shifts and rhumb-line bearings between present and 2021–2040, tested directional concentration with Rayleigh’s statistic, and mapped overlap, colonization, and extinction by overlaying present and future ranges. As a historical baseline, we analyzed 16,219,436 global herbarium records for 109,242 species across two periods (1970–2000; 2001–2024), and calculated 28-year centroid velocities and bearings, and compared realized versus projected rates using paired nonparametric tests. Predictors of velocity, colonization, and extinction (traits: height, evolutionary distinctiveness; biotic: ant richness(57); abiotic: soil N(58), soil organic carbon(59), elevation; anthropogenic: protected-area coverage, threat(60), distances) were centered and scaled and analyzed with linear mixed-effects models including clade/family and biome as random effects.

## Data, Materials, and Software Availability

The analyses were performed using open source and reproducible tools in R 4.5.1 (2025-06-13), “Great Square Root” using packages terra v.1.7-3, dismo v.1.3-14, rangeBuilder v.2.1, castor v.1.7.10, V.PhyloMaker2 v.O.1.0, spatialEco v.2.0-1, usdm v.2.1-6, MaxEnt v.3.4.3, phyloregion v.1.0.9, sf v.1.0-21, geosphere v.1.5-20, and lmerTest v.3.1-3.

## Acknowledgments

We are grateful for financial support from the U.S. National Science Foundation (awards 2345994 and 2416314), Alfred P. Sloan Foundation, and Stanford Woods Institute Big Ideas for Oceans. The authors acknowledge the Stanford Sherlock High Performance Computing Center (https://sherlock.stanford.edu) for providing HPC resources that have contributed to the research results reported within this paper.

